# The different brain areas occupied for integrating information of hierarchical linguistic units: a study based on EEG and TMS

**DOI:** 10.1101/2021.11.22.469495

**Authors:** Changfu Pei, Yuan Qiu, Fali Li, Xunan Huang, Yajing Si, Yuqin Li, Xiabing Zhang, Chunli Chen, Qiang Liu, Zehong Cao, Nai Ding, Shan Gao, Kimmo Alho, Dezhong Yao, Peng Xu

**Affiliations:** The Clinical Hospital of Chengdu Brain Science Institute, MOE Key Lab for Neuroinformation, University of Electronic Science and Technology of China, Chengdu, China; School of Life Science and Technology, Center for Information in BioMedicine, University of Electronic Science and Technology of China, Chengdu, China; School of Foreign Languages, University of Electronic Science and Technology of China, Sichuan, Chengdu, China; School of Psychology, Xinxiang Medical University, Xinxiang, China; Institute of Brain and Psychological Sciences, Sichuan Normal University, Sichuan, Sichuan, Chengdu, China; STEM, Mawson Lakes Campus, University of South Australia, Adelaide, Australia; College of Biomedical Engineering and Instrument Sciences, Key Laboratory for Biomedical Engineering of Ministry of Education, Zhejiang University, Hangzhou, China; Department of Psychology and Logopedics, University of Helsinki, Helsinki, Finland

## Abstract

Human linguistic units are hierarchical, and our brain responds differently when processing linguistic units during sentence comprehension, especially when the modality of the received signal is different (auditory, visual, or audio-visual). However, it is unclear how the brain processes and integrates language information at different linguistic units (words, phrases, and sentences) provided simultaneously in audio and visual modalities. To address the issue, we presented participants with sequences of short Chinese sentences through auditory or visual or combined audio- visual modalities, while electroencephalographic responses were recorded. With a frequency tagging approach, we analyzed the neural representations of basic linguistic units (i.e., characters/monosyllabic words) and higher-level linguistic structures (i.e., phrases and sentences) across the three modalities separately. We found that audio-visual integration occurs at all linguistic units, and the brain areas involved in the integration varied across different linguistic levels. In particular, the integration of sentences activated the local left prefrontal area. Therefore, we used continuous theta-burst stimulation (cTBS) to verify that the left prefrontal cortex plays a vital role in the audio-visual integration of sentence information. Our findings suggest the advantage of bimodal language comprehension at hierarchical stages in language-related information processing and provide evidence for the causal role of the left prefrontal regions in processing information of audio-visual sentences.

## Introduction

Language allows us to communicate ideas, feelings, and needs and is the primary mark that distinguishes us from other species [1, 2]. Since human language is generally multisensory, it is a significant challenge to clarify how continuous language is perceived and integrated to build the meaning of sentences. Language comprehension involves integrating multisensory information (typically from auditory and visual modalities) to access meaning. Multisensory integration, defined as brain reactivity in response to the combination of signals from different modalities, is dynamic and context-dependent [3, 4]. To understand continuous speech, listeners must construct a linguistic structure at different hierarchies, including words, phrases, and sentences. Understanding naturally connected sentences depend on interconnections between word, phrase, and sentence processing. The three levels of linguistic units are different in terms of their functions in communication. It is generally believed that ‘sentence’ is the basic unit for speech communication, while ‘phrase’ and ‘word’ are standby units of language communication [5, 6]. However, there isn’t any neuro processing evidence to support this theory. Thus, an intriguing question is whether the combined audio-visual presentation can enhance the information processing of different linguistic units in naturally connected sentences. An increasing body of studies has explored the mechanisms underlying audio- visual integration of letters and speech sounds [7–9] and consistently demonstrated the superiority of audio-visual integration over processing letters and speech sounds separate [10, 11]. For example, a magnetoencephalography (MEG) study showed enhanced brain activity predominantly in the right temporal-occipital-parietal junction and the left and right superior temporal sulci for audio-visual integration of phonemes and graphemes [12]. This integration resulted in the reduced response of audio-visual (AV) stimuli in comparison with summated responses to unimodal auditory (A) and visual (V) stimuli in this study (i.e., AV < sumAV), which was interpreted as suppressive interaction. Thus, audio-visual integration can also be examined by brain responses evoked by audio-visual stimuli with the sum of responses to unimodal stimuli. If auditorily and visually presented synchronous stimuli were processed independently, then the neural responses induced by an audio- visual stimulus should be approximately close to the sum of the responses to unimodal stimuli presented separately. However, if the bimodal response differs in a supra-additive or sub-additive manner from the sum of the two unimodal responses, this is attributed to the interaction between the two modalities that can integrate the information. Atteveldt et al. used functional magnetic resonance imaging (fMRI) to investigate the integration of letters and speech and found the simultaneous presentation of auditory and visual stimuli modulated related activity in the superior temporal sulcus [7].

Since most studies concerning audio-visual integration have focused on the single letter-speech sound mapping in alphabetic languages, audio-visual integration on sentence-level has remained unveiled. In essence, the information of language is mostly conveyed at sentence-level, and the brain utilizes the more complicated scheme to process the language-related information in sentence-level compared to the other two lower levels. This research modified the experimental paradigm of Ding et al.[13] to study the audio-visual integration mechanism of different language units. To understand connected language, however, humans have to learn to construct a hierarchy of linguistic structures, including words/syllables, phrases, and sentences [14]. Cortical activity is synchronized with the acoustic features of speech approximately at the syllabic rate, which provides an initial timescale for speech processing, as well as for possibilities to explore its potential mechanism [15–19]. Sheng et al. compared neural activity synchronized with syllabic, phrasal, and sentential linguistic units in the frequency domain [20]. The superior temporal gyrus was found to be involved in the processing of the three linguistic units, while the activity in the motor cortex was associated with the processing of the rhythm of monosyllabic words, and both the left anterior temporal cortex and left inferior frontal gyrus were involved in the processing of phrases or sentences [13, 20].

Chinese is a logographic language comprising both auditory (syllabic) and visual (graphemic) characters. When perceiving Chinese characters, Chinese speakers usually integrate multisensorial information, that is, visual and auditory features, and construct a hierarchy of different linguistic structures, including words, phrases, and sentences [12, 13, 20]. Yet, it remains an open question whether audio- visual integration is superior over single auditory or visual processing when the brain simultaneously handles the three linguistic structures at different timescales [21–24]. Therefore, in the present study, we hypothesized that audio-visual integration outperforms unimodal processing in logographic languages. Thereafter, the corresponding electroencephalographic responses at different timescales [25, 26] were collected with electroencephalography (EEG) in a hierarchical linguistic sequence paradigm (Fig 1). To identify the potential differences, we further compared participants’ behavior and multiple aspects of brain responses, including spectral (frequency) [13, 20], time (event-related potentials (ERPs)) domains, as well as the brain networks involved, between the auditory, visual, and audio-visual conditions (Fig 2). Finally, based on the results, the related brain areas specifically for language information integration will be revealed. To validate the role of those brain areas for information integration, we enrolled another independent group to attend our second experiment consisting of sham stimulation and cTBS (Fig 3). Unlike the actual stimulation exerted on the brain site by cTBS, sham stimulation only places the transcranial magnetic stimulation (TMS) device on the brain site and does not give the proper stimulation. Then, after the stimulation, the participants attend the experiment following the protocol in Experiment 1, during which EEG and behavior responses are recorded, aiming to probe whether the participant’s capability to understand the language will be influenced when the critical brain areas for language information integration is modulated.

**Fig 1.**
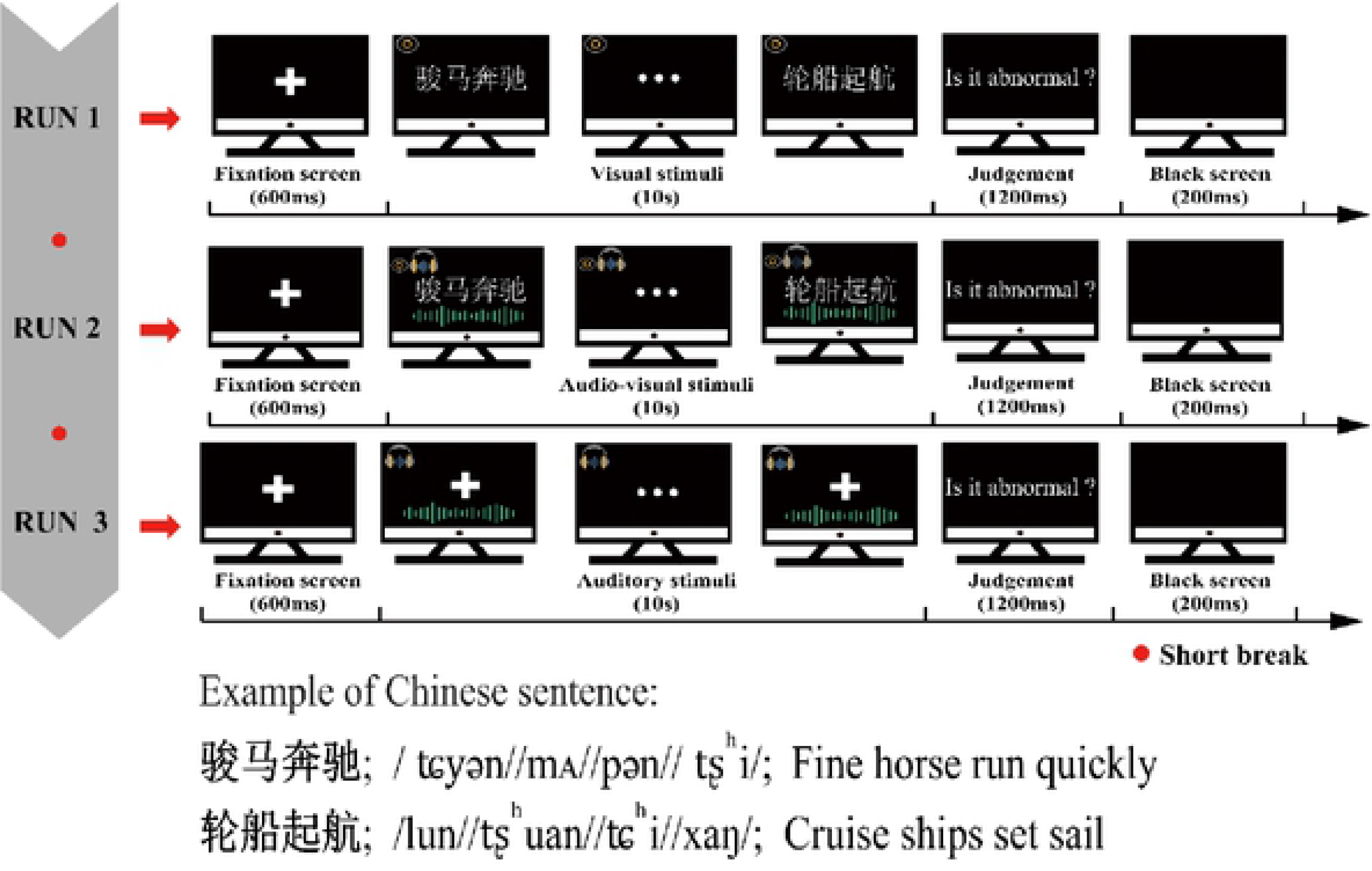
Schematic of the experimental design. A schematic standard (‘normal’) trial of the visual (top), audio-visual (middle), and auditory (bottom) condition consisting of a 600-ms cue, 10-s stimulus sequence, 1200-ms judgment, and 200-ms break. Below the Chinese stimulus examples, both English translation and phonological transcription in the International Phonetic Alphabet (IPA) are presented.

**Fig 2.**
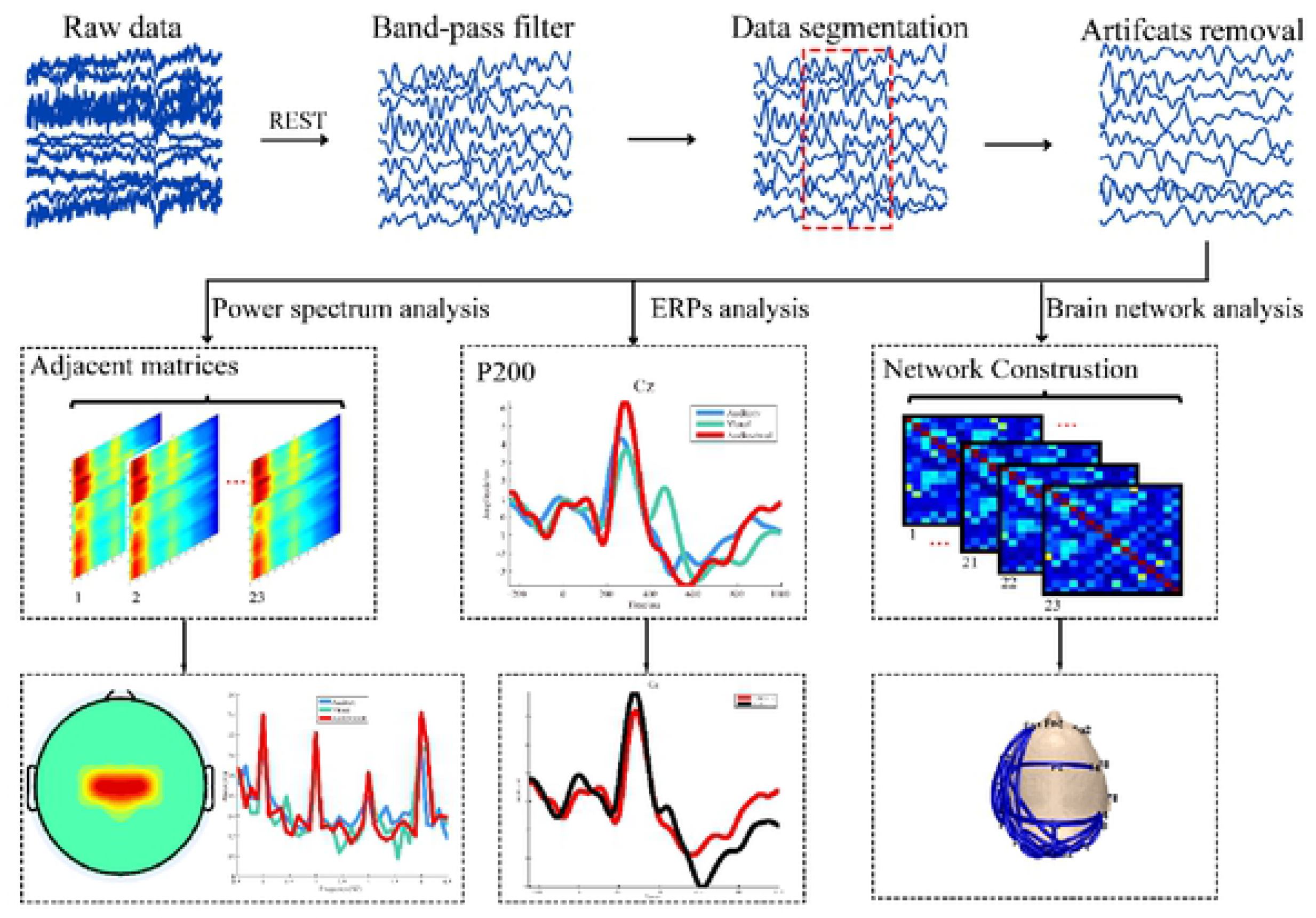
The analysis procedure for EEG data.

**Fig 3.**
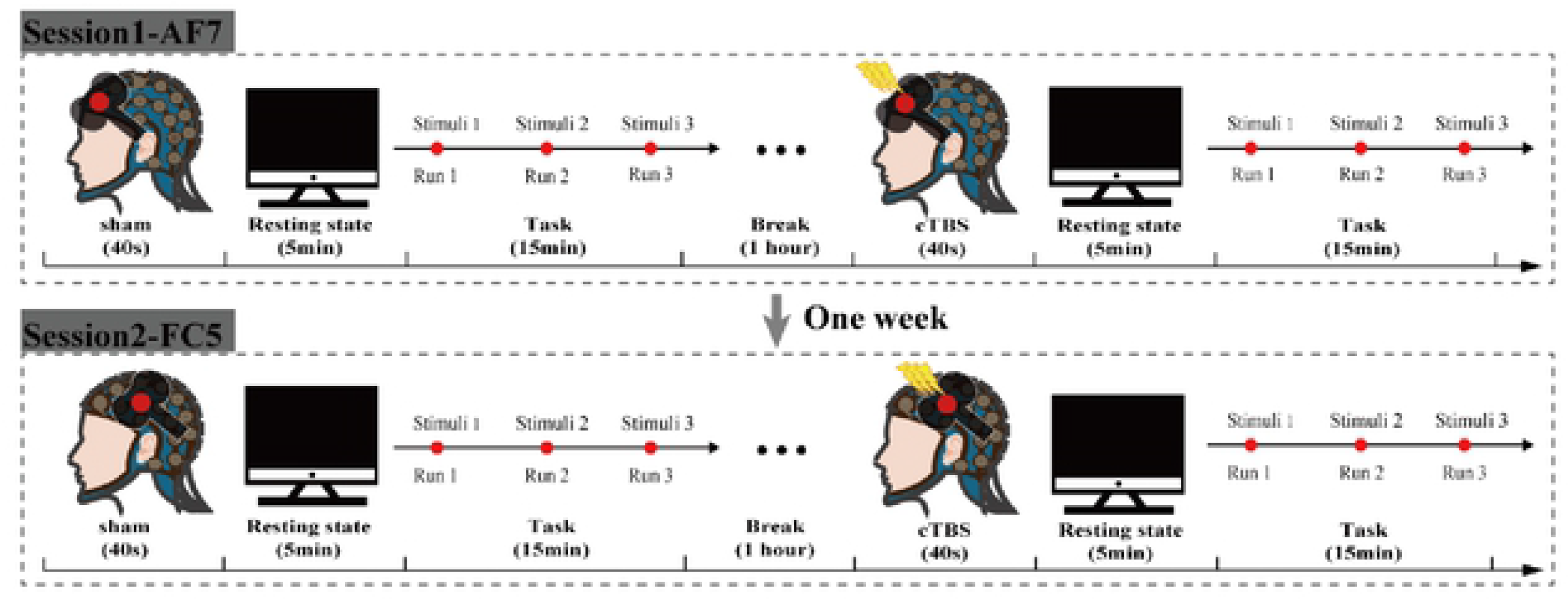
A schematic description of the two site sessions. The procedure was similar for both sessions, with the only difference that one of them was AF7 site and the other one was FC5 site.

## Results

### Experiment 1

#### Behavioral differences

As illustrated in Fig 4 (left panel), the mean (± standard deviation) of accuracy (ACC) was 97.0 ± 3.0% in the auditory condition, 95.7 ± 4.3% in the visual condition, and 98.2 ± 2.6% in the audio-visual condition. The mean (± standard deviation) of reaction time (RT) was 538.4 ± 158.8 ms in the auditory condition, 587.9 ± 150.9 ms in the visual condition, 454.7 ± 119.7 ms in the audio-visual condition, as shown in the right panel of Fig 4. The repeated-measure ANOVAs identified a significant main effect of modality (ACC: *F*_(2,44)_ = 5.85, *p* = 0.01; RT: *F*_(2,44)_ = 7.67, *p* = 0.006). The post-hoc tests further showed that ACC was significantly higher in the audio-visual condition than in the visual (*F*_(1,22)_ = 10.14, *p* = 0.004) and auditory (*F*_(1,22)_ = 5.28, *p* = 0.03) conditions. RT was significantly shorter in the audio-visual condition than in the visual (*F*_(1,22)_ = 21.95, *p* < 0.0005) and auditory (*F*_(1,22)_ = 10.02 , *p* = 0.004) conditions.

**Fig 4.**
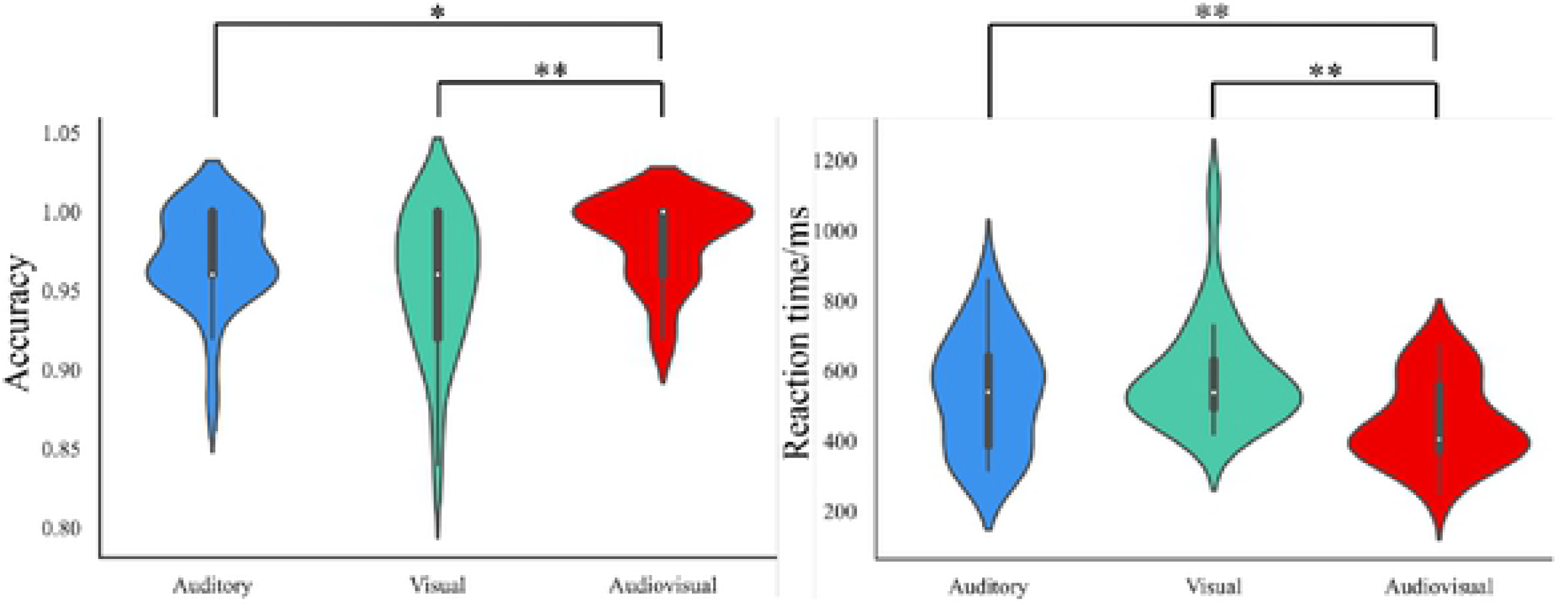
Behavioral performance. The mean ACC and RT in detecting “abnormal” (outlier) trials in the auditory, visual, and audio-visual conditions, shown in the form of violin plots. White dots represent the median and the first and third quartiles are identified by the bottom and top of the bold vertical lines, respectively. The bottom and top of the thin vertical line represent the lower and upper adjacent values, respectively (* *p* < 0.05, ** *p* < 0.01, Bonferroni-corrected).

#### Power spectrum

The monosyllabic words, phrases, and sentences were presented every 0.25 s, 0.5 s, and 1 s, respectively. As illustrated in Fig 5, the frequency of brain responses to each linguistic unit was synchronously tagged, that is, 4 Hz for words, 2 Hz for phrases, and 1 Hz for sentences. Concerning the power spectrum at the electrode Cz, the repeated-measure analyses of variance (ANOVA) revealed a significant main effect of modality (*F*_(2,44)_ = 8.678, *p* = 0.001) such that the power was enhanced in the audio-visual (23.55 ±15.72 dB) condition compared with either the auditory (20.55 ±11.08 dB, *p* = 0.004, *F*_(1,22)_ = 10.65) or the visual (21.37 ± 14.26 dB, *p* = 0.001, *F*_(1,22)_ = 14.21) conditions. A significant main effect was found for linguistic units (*F*_(2,44)_ = 3.894, *p* = 0.04). The power spectrum was stronger for sentential linguistic units than other linguistic structures (auditory: 21.07 ± 19.56 dB; visual: 23.04 ± 23.69 dB; audio-visual 23.98 ± 24.80 dB). However, the pairwise comparisons showed no significant differences. No significant interaction of modality and type of linguistic unit was observed.

**Fig 5.**
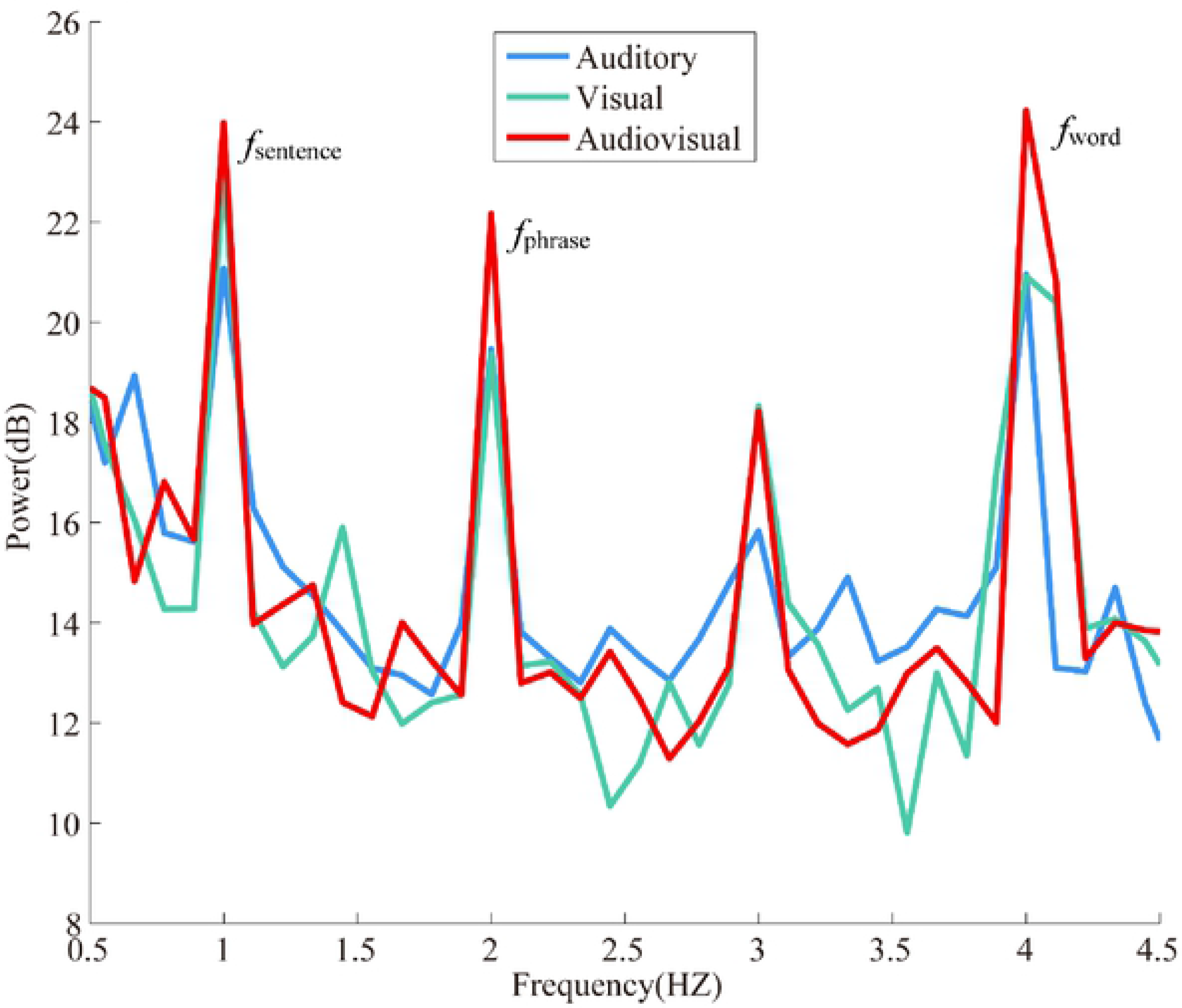
Neural tracking of hierarchical linguistic structures under different modalities. The grand-averaged power spectrum at the electrode Cz in the auditory, visual, and audio-visual conditions. Neural tracking of syllabic, phrasal and sentential rhythms was reflected by spectral peaks at corresponding frequencies.

As seen in Fig 6, there were significant scalp topography differences in the power for the three hierarchical units among auditory, visual, and audio-visual conditions. For word processing, significant differences between the audio-visual and visual conditions (Fig 6A, left panel) and the audio-visual and auditory conditions (Fig 6A, right panel) were observed over the central midline areas. For phrasal processing, the left frontal and the bilateral parietal regions showed significantly more vigorous activity in the audio-visual condition than in the visual condition (Fig 6B, left panel). In contrast, significant differences were found over the left parietal and the frontal areas between the audio-visual and auditory conditions (Fig 6B, right panel). For sentential processing, left frontal areas showed stronger activity in the audio-visual condition than in the visual (Fig 6C, left panel) or auditory condition (Fig 6C, right panel).

**Fig 6.**
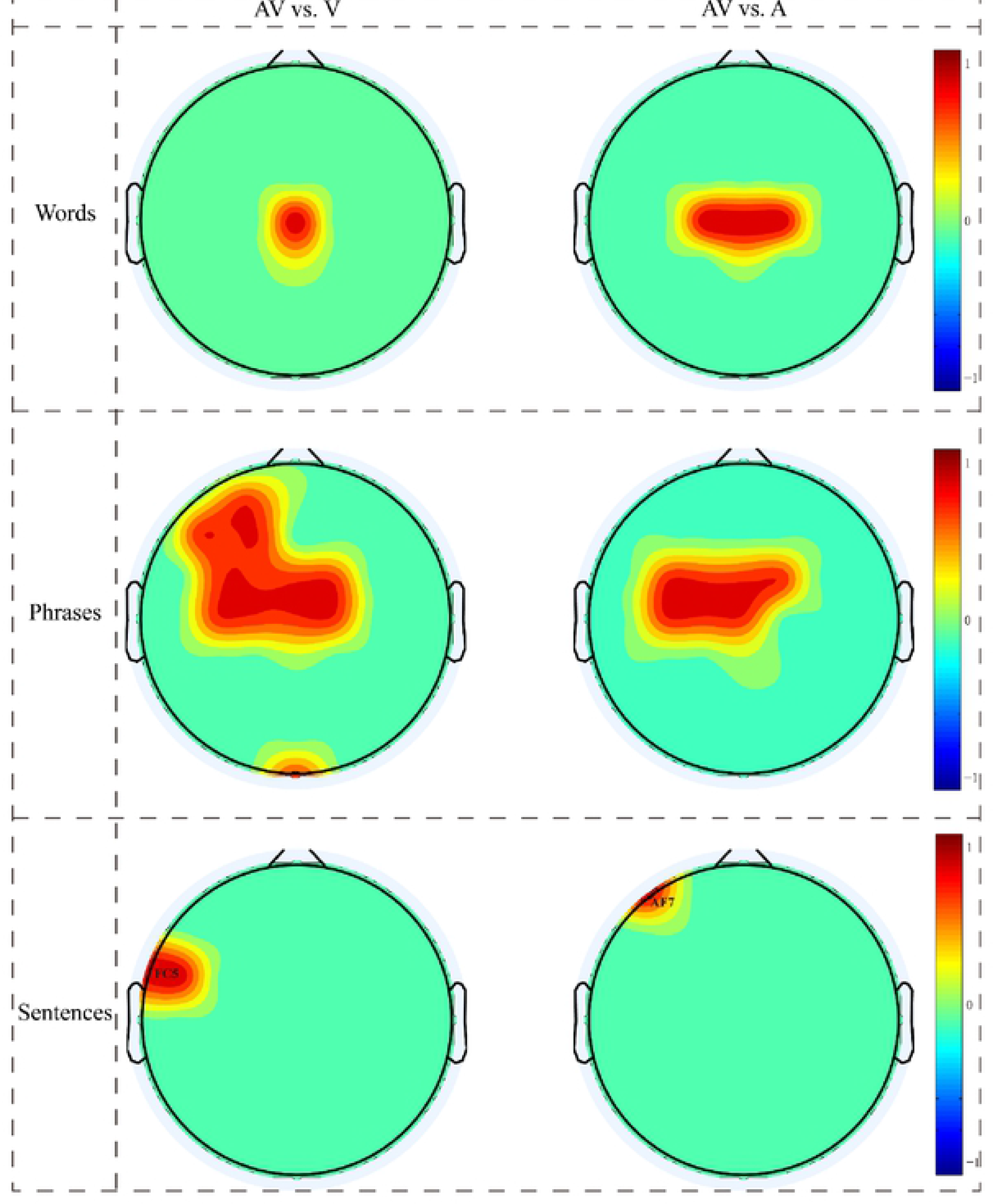
Topography differences of power among the three conditions for different linguistic units. Red indicates significantly increased activation (*p* < 0.01 paired *t*-test, Bonferroni-corrected); blue, not seen in the scalp illustrations, would indicate significantly decreased activation. auditory, A; visual, V; audio-visual, AV.

#### ERPs results

As Fig 7 depicts, the P200 amplitudes were significantly different among modalities (*F*_(2,44)_ = 10.17, *p* < 0.001). Pairwise comparisons revealed that the P200 was enhanced in the audio-visual (4.77 ± 0.37 μV) condition compared with either the auditory (3.51 ± 0.46 μV, *p* < 0.002) or the visual (2.86 ± 0.36 μV, *p* < 0.001) conditions.

**Fig 7.**
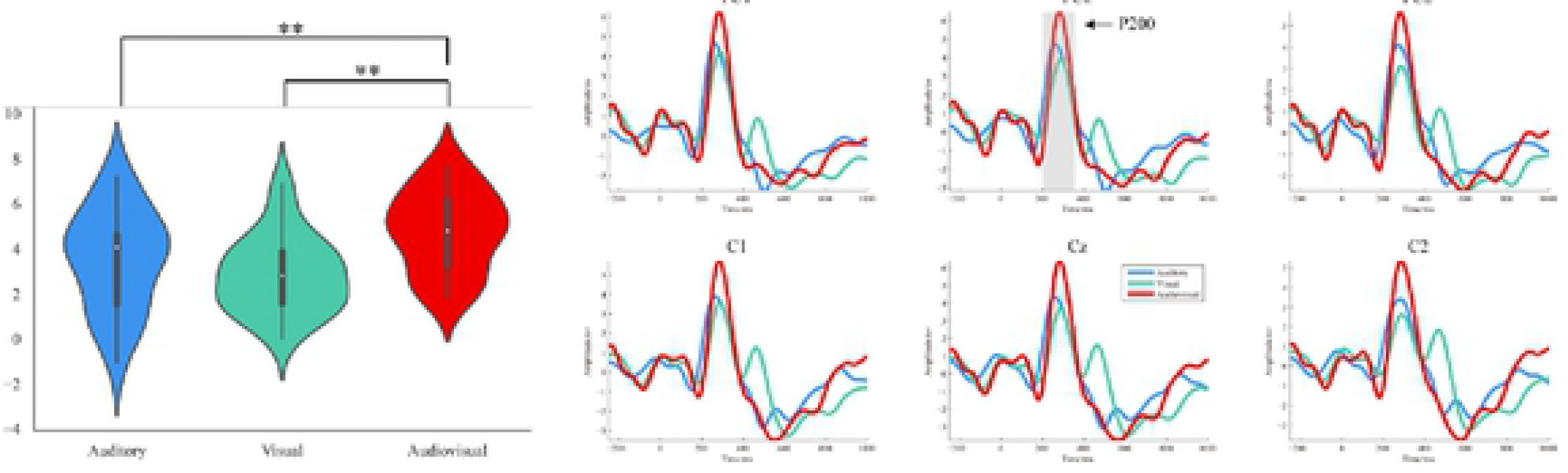
Grand-averaged ERPs elicited by sentences under the different modalities. The left panel displays that the mean amplitude of P200 over six electrodes (FC1, FCz, FC2, C1, Cz, and C2) in the auditory, visual, and audio-visual conditions, shown in the form of violin plots (* *p* < 0.05, ** *p* < 0.01, Bonferroni-corrected). White dots represent the median, and the first and third quartiles are identified by the bottom and top of the bold vertical lines, respectively. The bottom and top of the thin vertical line represent the lower and upper adjacent values, respectively. The right panel displays brain potential variations over the six electrodes expected to ideally capture the P200. The shaded area highlighting the P200 are approximative; for accurate analysis time-windows, see Materials and Methods.

Fig 8 compares the sumAV with the P200 in response to audio-visual stimuli, showing reduced P200 during audio-visual integration (*t* = -2.72, *p* < 0.012). In terms of the scalp distribution of AV minus sumAV difference for the P200, in the left frontal area and central scalp region, the amplitude of P200 was smaller under AV than sumAV (see the right panel of Fig 8).

**Fig 8.**
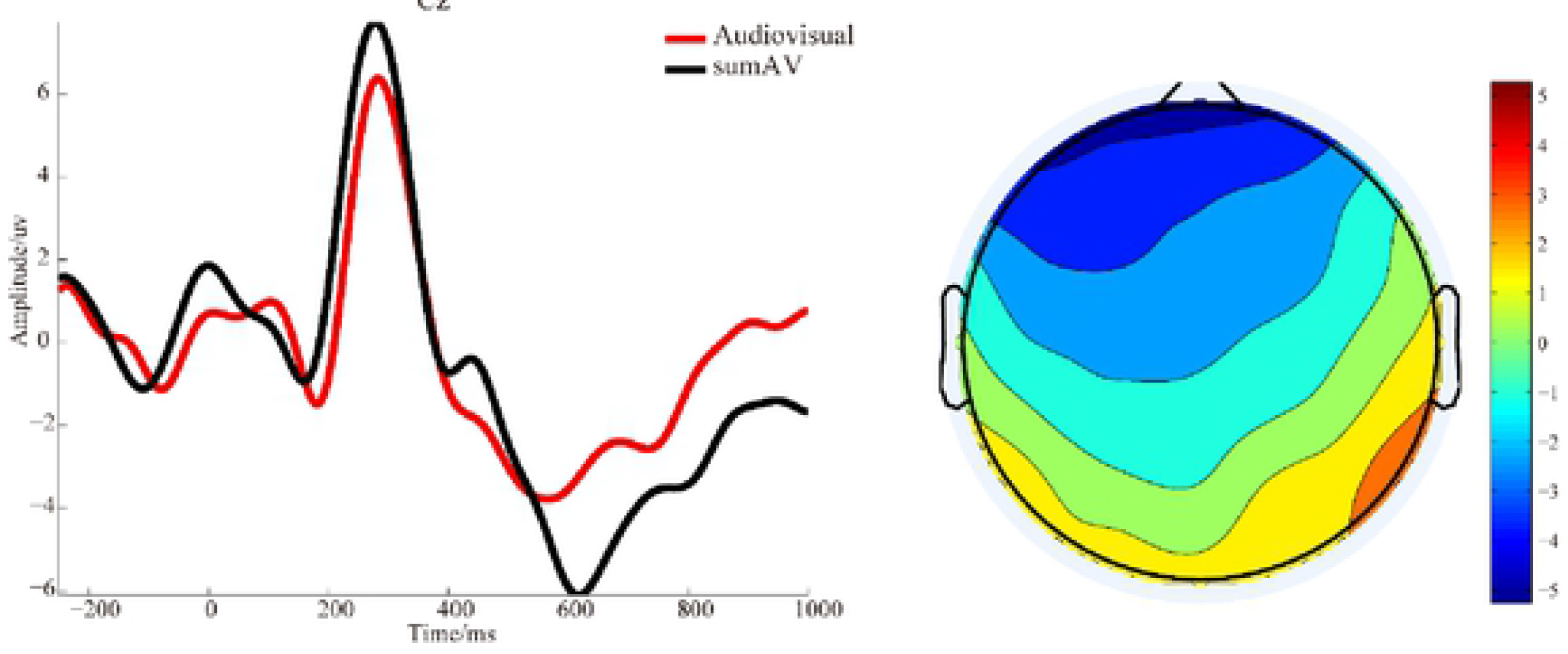
The comparison between ERPs responses to audio-visual sentences and the sum of ERPs in response to the two unimodal stimuli. The left panel displays the grand-averaged ERPs at the electrode Cz. The right panel displays the topographic AV-sumAV map for the P200.

#### Patterns of brain network in different conditions

The identified differences of the network architectures between the audio-visual condition and the unimodal visual condition (*p* < 0.01, paired *t*-tests; Fig 9) showed enhanced linkages in the audio-visual condition over widely distributed scalp areas. The linkage enhancements were left-hemisphere dominant, especially in relation to the unimodal auditory condition. No decreased linkages in the audio-visual condition compared with the unimodal condition and no significant linkage differences between the visual and auditory conditions were observed.

**Fig 9.**
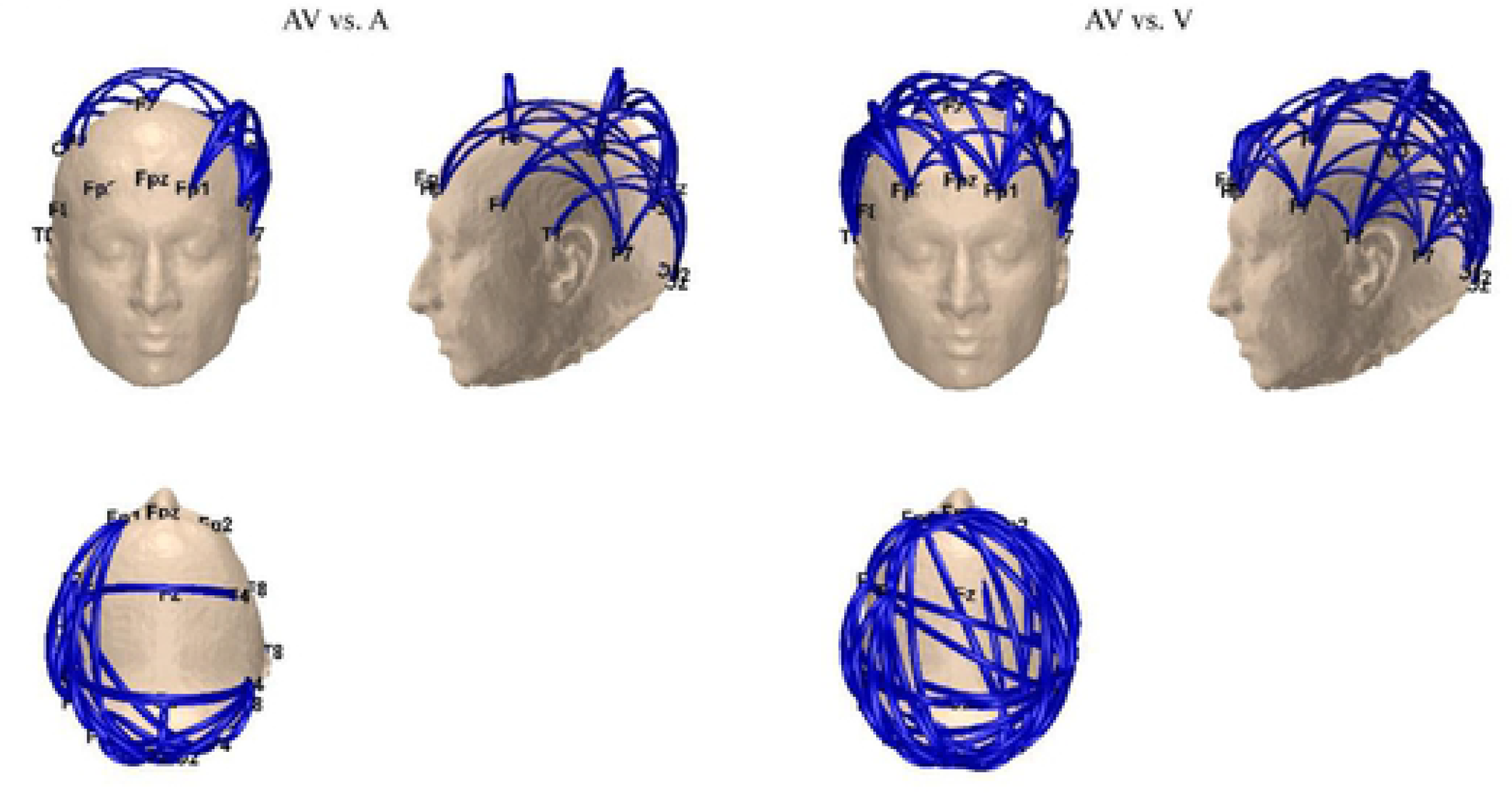
The difference network topologies between the bimodal and unimodal conditions. The blue lines denote the edges with statistically stronger (*p* < 0.01; paired *t*-tests) linkages in the audio- visual condition than in the visual or auditory condition.

### Experiment 2

Experiment 1 identified two nodes (i.e., AF7 and FC5 revealed by power spectrum analysis) that played a critical role in the audio-visual integration of sentential structure. To validate whether behavioral and electrophysiological responses would change when the hub nodes were modulated, we conducted Experiment 2 where TMS was employed to regulate the activity at the concerned nodes. Previous evidence had shown that when TMS was applied to AF7, the stimulation could be transferred to modulate the activity at the left prefrontal region [39]. There is also research reporting that the function in the precentral gyrus was affected by giving TMS application to FC5 [40]. Therefore, we expected the changes both in their behavior and EEG responses to an audio-visual stimulus will be observed when the activities of the two nodes are suppressed by TMS, which is mainly attributed to the disturbance of the audio-visual information integration for sentential structure.

#### Behavioral differences

In term of ACC, the interaction between TMS condition and modality conditions was significant (*F*_(2,28)_ = 3.79, *p* = 0.035; S1 Fig). The pairwise comparisons revealed that in the audio-visual and auditory condition ACC displayed a reduction following cTBS as compared to sham stimulation (audio-visual: *F*_(2,28)_ = 4.42, *p* = 0.054; auditory: *F*_(2,28)_ = 4.05, *p* = 0.06) while in the visual conditions no significant differences were observed between cTBS and sham stimulation (*F*_(2,28)_ = 0.88, *p* = 0.36). RT did not show a significant TMS-modality interaction.

#### Power spectrum

A significant three-way interaction was observed among TMS condition, power spectrum frequency, and modality condition (*F*_(4,56)_ = 3.35, *p* = 0.019; S2 Fig). The post-hoc comparisons revealed that in the audio-visual stimulation condition, power spectrum strength was significantly lower following cTBS as compared to sham stimulation (*F*_(4,56)_ = 10.15, *p* = 0.01) while in the unimodal conditions no significant differences were identified between cTBS and sham stimulation (auditory: *F*_(4,56)_ = 1.11, *p* = 0.32; visual: *F*_(4,56)_ = 0.07, *p* = 0.79). There were no other significant differences in other TMS – related effect.

#### ERPs results

The ANOVA on P200 amplitudes showed that a marginally significant interaction between TMS and modality conditions (*F*_(2,28)_ = 3.08, *p* = 0.068; S3 Fig) such that in the audio-visual stimulation condition the P200 was significantly smaller following cTBS as compared to sham stimulation (*F*_(2,28)_ = 7.99, *p* = 0.02) whereas in response to unimodal stimuli no significant differences were obtained between cTBS and sham stimulation (auditory: *F*_(2,28)_ = 1.77, *p* = 0.21; visual: *F*_(2,28)_ = 0.41, *p* = 0.54). S3 Fig depicts similar TMS-induced changes on AF7 and FC5 in response to audio- visual stimuli. In addition, the main effect of TMS was significant (*F*_(1,14)_ = 7.70, *p* = 0.02) with P200 amplitudes being reduced following cTBS relative to sham stimulation regardless of site and modality. No other TMS-related effects were observed.

## Discussion

Is the combination of auditory and visual inputs more conducive to language processing than unimodal inputs alone? In the present study, we addressed the questions in an experiment combining auditory (syllabic) and visual (graphemic) presentations of different hierarchical linguistic units of Chinese, given that understanding how the different levels of linguistic units are represented in the brain is the key to clarify the neural basis of language comprehension [13]. To these ends, first, the present study investigated the possible advantages of audio-visual integration for language-related information processing, and also probed the specific brain areas involved in this integration for the different hierarchical linguistic units, in terms of behavioral performance, EEG-based power spectrum, ERPs, and functional brain networks. Second, TMS was used to suppress modulate the activity of two hub nodes (AF7 and FC5) identified in the power spectrum analysis, aiming to validate the role of these two brain sites during the audio-visual integration of sentential structures.

In Experiment 1, participants responded more accurately and faster to audio- visual stimuli than they did in either of the unimodal conditions, indicating that spoken syllables and their orthographic information were successfully integrated and facilitated linguistic processing in the audio-visual condition. Motivated by the close relationship between electrophysiological activity and behavior in previous studies [25, 41], we further probed how syllabic and glyphic information is integrated into the brain.

We observed power spectrum peaks at 1 Hz, 2 Hz, and 4 Hz, consistent with previous studies [13, 20] and correspond to the rates of the sentence, phrase, and monosyllabic word presentations, respectively. While previous studies found information integration of language for monosyllabic words and phrases [21], audio- visual integration on the sentence level has remained unveiled. The current results provided evidence in this aspect, showing that the most robust responses at 1 Hz, 2 Hz, and 4 Hz for the audio-visual condition, suggesting that audio-visual integration not only exists for syllables (monosyllabic words) and phrases but also for the higher sentence-level linguist processing.

Though information integration was observed for all the three linguistic units, the scalp topography results showed that the different linguistic units involved different brain areas for the information integration. Specifically, in the audio-visual condition as compared to the unimodal condition, the processing of words led to more significantly stronger activation in the parietal areas. This is in line with a previous MEG study in Finnish school children that emphasizes the crucial role of the parietal-temporal cortex in the early phase of reading [9]. The parietal areas may be involved in early audio-visual integration [12, 42, 43]. For the phrasal processing, the audio- visual integration recruited the left prefrontal and bilateral parietal areas. The literature has shown that the left prefrontal area is primarily engaged in the processing of basic syntactic/semantic combinations [43]. Our result further verified the involvement of syntactic processing in phrases under the audio-visual modality. The topological activation with regard to sentences showed stronger responses in the left prefrontal area under the audio-visual relative to unimodal conditions. The region has not been ever found to be engaged in audio-visual integration concerning other types of linguistic units, including the letter-speech-related bimodal integration, which may be the specific brain area for audio-visual information integration of language [44–48]. The area with increased activation is part of Broca’s area. This is also consistent with the previous finding that the syntactic and semantic processing of sentences is associated with Broca’s area [23, 49–51]. Taken together, while audio-visual integration occurs at all the linguistic levels, the brain areas involved in the integration were different across the levels. Intriguingly, the processing of words and phrases showed some overlap in terms of brain activation, while sentential processing is rather different compared with them.

The results also indicate different degrees of hemispheric lateralization when integrating information in different hierarchical linguistic units. In detail, the integration of basic syllabic processing at the pre/post-central areas showed no hemispherical lateralization, potentially a motor-sensory network, which is consistent with previous studies reporting that the audio-visual integration of syllables is no significant hemispheric laterality [12]. However, for higher-level linguistic units, such as phrases and sentences, stronger activation due to information integration was observed in the left frontal and parietal areas, which is in line with previous findings that syntactic and semantic information is mainly processed in the left prefrontal and the left parietal areas [50, 52].

In terms of ERPs, the audio-visual condition showed larger amplitudes of the P200 as compared to the unimodal conditions. The P200 component, in general, is known to reflect the allocation of attention [53, 54]. When the brain processes audio- visual stimuli, both visual and auditory attention resources are needed [42]. Therefore, more attention may be allocated to the audio-visual modality relative to unimodal stimuli here, leading to enhanced amplitudes of the P200. The P200 has also been shown to be sensitive to semantic priming [55, 56]. The semantic priming may indeed require the conscious linking of related representations. Such a mechanism would be superfluous for cross-modal repetition priming since the visual glyphs, and auditory speech presumably has the same semantic representation. It is likely that audio-visual integration in language processing requires more cognitive resources than unimodal processing to facilitate semantic/syntactic understanding [42, 57, 58] and thus produced larger P200 amplitudes. In agreement with previous evidence for suppressive interaction in the auditory and visual processing of audio- visual stimuli [12]. the P200 responses to audio-visual stimuli in the present study were smaller than the sum of ERPs elicited by both types of unimodal stimuli (sumAV > AV). This sub-additive effect may reflect the facilitation of auditory and visual processing due to the audio-visual presentation of the same stimulus. In the topographic AV-sumAV map for the P200, the AV < sumAV areas were predominantly over the left frontal and central regions. This left hemisphere dominance in scalp distribution might be related to linguistic audio-visual integration. When children learn to read, written and spoken words are often presented together and neural pathways that enable them to memorize and retrieve audio-visual associations are formed. Consequently, the audio-visual suppression effect can be interpreted as optimization of neural networks during learning [12].

In addition to local brain activity reflected by power spectrum responses, we conducted the brain network analysis, which could structure how the information is propagated among the related brain areas. Consistent with increased activities for audio-visual stimuli, we identified increased network patterns in the audio-visual relative to unimodal modality, namely, simultaneous integration of visual and auditory information recruited more brain sources responsible for language-related information processing. Another aspect revealed is that the enhanced linkages under audio-visual conditions are exhibited with the lateralization in the left hemisphere, which is consistent with the observed lateralization for the power spectrum. Despite increased network patterns for the audio-visual stimuli as compared to both visual and auditory stimuli, the increased network patterns differed when comparing the bimodal modality with either of the two single modalities. Compared with the differences between the audio-visual and auditory stimuli, the differences between the audio-visual and visual stimuli were more pronounced. This difference suggests that when language information is visually represented, the linguistic information is not efficiently processed in the brain that evoked a less efficient network, while the audio stimulus evoked a more efficient network for information processing. Behaviorally, the accuracy in visual stimulus presentation was associated with the lowest accuracy, which may indicate that the visual stimulus is not competitive for language processing. In the visual modality, the pronunciation of words is activated first from visual glyphs and then from the pronunciation of words to the meaning, and therefore the visual modality is less efficient than the auditory modality, which allows the linguistic information to be processed directly from pronunciation to meaning [59]. Moreover, the auditory modality evoked the stronger P200 compared to the visual modality as shown in Fig 7, which suggests that the auditory modality may recruit more brain resources and thus promote processing. This superiority of the auditory modality for language to the visual modality for language processing may be attributed to the usual way of language acquisition that an infant initially receives the language from hearing [60], and the auditory properties of language during the communications [61].

The present study (Experiment 2) investigated the causal role of the left prefrontal regions in the audio-visual integration in sentence processing. Consistent with our hypothesis, the inhibition of the left prefrontal region by cTBS changed the behavioral and electrophysiological responses. When cortical excitability of these areas was decreased via TMS, a significant increase in RT and a decrease in ACC were observed, indicating a reduction in audio-visual integration. Following the cTBS intervention (over the electrodes AF7 and FC5), compared with the sham condition, the P200 and power spectrum were significantly reduced. Given that the P200 has been associated with the allocation of attention resources and semantic priming [53, 55], smaller P200 amplitudes following cTBS may suggest that the stimulation cuts down on cognitive resources that the audio-visual integration in language processing demands and thus undermines semantic judgment. When TMS was applied to AF7, compared with the sham condition, there was a significant drop of the sentence-level power spectrum (1Hz) in the audio-visual modality. Although the power spectrum of word-level (4Hz) and phrase-level (2Hz) decreased, there was no significant TMS effect under the audio-visual condition. Moreover, for the electrodes FC5, there was no statistical difference in sentence-level (1Hz) power spectrum under audio-visual modality compared to the sham condition. These results confirm that the left prefrontal cortex may also be responsible for audio-visual integration at the sentence level.

## Conclusion

The ability to read is a major landmark process in human cognitive development. Cognitive scientists and psycholinguists have maintained that learning to read requires skills in orthographic, phonological, and semantic facets of printed words. Therefore, learning to read involves the integration of multisensory (syllabic and graphemic) information to access meaning. When children start understanding written language, they typically learn to associate the sounds of their spoken language with unfamiliar characters in the logographic language and finally access the meaning of visual glyphs. The results of Experiment 1 based on multiple measurements consistently identified the superiority of audio-visual integration over a single auditory or visual modality in hierarchical linguistic units. Higher ACC and shorter RT, concurrent with enhanced EEG power spectra and their topological distributions, larger P200 amplitudes, the difference of sumAV and AV for P200, and more network linkages, suggested that the audio-visual modality could facilitate the semantic/syntactic understanding of Chinese. In Experiment 2, after TMS was used to suppress the activity of the nodes obtained by power spectrum analysis, we observed the significant changes in behavioral responses and electrophysiological indices for audio-visual integration. Overall, our results suggest that audio-visual integration as compared to unimodality produces an advantage in processing hierarchical linguistic units in Chinese via the left prefrontal cortex and that listening and reading at the same time may be an effective way to learn and understand language, especially hieroglyphs like Chinese. Given that learning to integrate visual glyph and pronunciation is crucial to acquire the ability to read a language, our results may have important implications for the acquisition of reading skills.

## Materials and Methods

### Experiment 1

#### Participants

The Institution Research Ethics Board of the University of Electronic Science and Technology of China (UESTC) approved the experiment. We recruited 23 healthy, right-handed postgraduates (13 males, 10 females; age 24.09 ± 2.48 years) from the student population at the UESTC. They were all right-handed native Chinese speakers and had normal hearing and normal or corrected to normal visual acuity. Participants had never used any psychoactive medication, and none had a personal or family history of psychiatric or neurological illnesses. Before the experiment, written informed consent was obtained from all participants after they fully understood the procedure.

#### Stimuli

The stimuli in the experiment included 50 short sentences composed of four Chinese monosyllabic words, with the first two syllables constituting a noun phrase and the last two syllables constituting a verb phrase [13]. In these sentences, noun and verb phrases were compatible. A ‘normal’ (standard) trial consisted of ten meaningful sentences. An abnormal trial consisted of eight meaningful sentences and two nonsense sentences derived from the meaningful sentences by reversing their subjects and predicates. For example, based on two compatible sentences, 轮船起航 (cruise ships set sail) and 青草发芽 (green grass grew bud), two nonsense sentences were made: 轮船发芽 (cruise ships grew bud) and 青草起航 (green grass set sail). All sentences were provided in three modalities: auditory, visual, and audio-visual. The duration of each syllable in the auditory sentences was adjusted to 250 ms and the gap between adjacent syllables was removed to avoid a potential contribution of speech rate or any other prosodic cue to linguistic structure building. Thus, the presentation rate was 4 Hz, which is close to the mean syllable rate in natural speech across languages [13, 20, 27]. The visual sentences consisted of four sequentially presented Chinese characters. The stimuli of spoken syllables and corresponding characters in the audio-visual condition were synchronously delivered. The speech inputs were delivered binaurally via headphones, and their intensity was about 65 dB SPL. The written characters (size: ca. 25.31 mm × 25.31 mm) were delivered in white font on a black computer screen at a distance of about 70 cm in front of the participant.

#### Experimental procedures

Each participant was presented with sentences in the three modality conditions (i.e., auditory, visual, or audio-visual). In each condition, Chinese monosyllabic words were presented in sequences in random order or an order forming the two-phrase sentences. The sentences of one modality were presented in one run, resulting in 3 runs. Each run consists of 25 trials including 20 standard trials where ten compatible sentences were presented without acoustic gaps and 5 outlier trials where two nonsense sentences were presented among the ten sentences. The order of sentences and trials within a run was randomized. Each trial lasted a period of 12 s (Fig 1), starting with the presentation of a fixation cross for 600 ms. Following the fixation, ten two-phrase sentences were presented, each two-phrase sentence lasting for 1000 ms. After the presentation of sentences, participants were required to judge whether the trial was “normal” (standard) or not. If the trial was “abnormal” (outlier), participants should press key “1” on a keyboard within 1200 ms, while if the trial is ‘normal’, participants need not press any button. After a 200 ms blank period, the next trial was initiated. Before the experiment, all participants were required to try a preliminary round to ensure that they understood the rules of the experiment. There is a 3-minutes interval between two consecutive runs for rest, and the order of runs (i.e., auditory, visual, or audio-visual) are randomly presented for each participant.

#### Task performance

ACC and RT per condition (auditory, visual, and audio-visual) was recorded. Both button presses to abnormal/outlier trials and omissions of button presses to normal/standard trials were regarded as correct responses.

#### Data acquisition

Participants were seated comfortably in an electrically shielded, dimly lit room. EEG data were recorded with 64 Ag/AgCl electrodes (ANT Neuro, Berlin, Germany), and all electrodes were positioned according to the extended 10-20 international electrode placement system (ASA-Lab Amplifier, eemagine Medical Imaging Solutions GmbH, Berlin, Germany). The online sampling rate was 500 Hz, and the data were band-pass filtered at 0.01–100 Hz. The electrodes CPz and AFz served as the reference and ground, respectively. To monitor eye movements, an electrooculogram was recorded using an additional electrode positioned above the left eye. During the entire task, the impedances of all electrodes were kept below 5 kΩ.

#### Data analysis

A series of procedures consisting of pre-processing, power spectrum calculation, ERPs extraction, and network analysis were implemented (Fig 2).

#### Pre-processing

Reference electrode standardization technique (REST) referencing (http://www.neuro.uestc.edu.cn/rest/) [28, 29], offline band-pass filtering, data segmenting, and artifact-trial removal were included to pre-process the recorded EEG. Concerning the power spectrum analysis, [0.1, 10] Hz offline band-pass filtering, [-800, 0] ms baseline correction, and artifact-trial removal with a threshold of ± 75 μV were used. For each trial, the stimulus period was defined by ignoring the first second of the stimulus to avoid the transient response [20]. Thus, the length of the segmented data was 9 s.

#### Power spectrum

The direct Fourier transform was applied to each artifact-free trial to calculate the power spectra per condition. Thereafter, for each modality, the corresponding power spectra were averaged across all trials to acquire the final response power [20].

#### ERPs analysis

Following pre-processing (Fig 2), epochs ranging from -250 to 1000 ms after the onset of the final word in a sentence were averaged for each modality. Peak detection was performed automatically, time-locked to the latency of the peak at the electrode of maximal amplitude on the grand-average ERPs. Temporal windows for peak detection were determined based on variations of the global field power measured across the scalp [30]. The P200 was defined as the mean amplitude in the 250–350 ms time window following word onset at six electrodes over the frontocentral and central areas (i.e., FC1, FCz, FC2, Cz, C1, and C2) where the P200 is classically found and displays maximal sensitivity [31, 32]. ERPs to separately delivered auditory (A) and visual (V) stimuli were summed, and this sum (sumAV) was compared with audio-visual stimuli ERPs (AV) to investigate whether audio- visual integration could be reflected by the ERPs of AV stimuli i.e., sub-additivity (AV < sumAV) for information integration.

#### EEG network

We adopted the phase-locking value (PLV) [33, 34] that can capture the non-linear phase synchronization between paired nodes to construct language-related network. To reduce the volume conduction [35, 36], we sparse 21 canonical electrodes as network nodes to construct the networks. To estimate the corresponding instantaneous phases, i.e., *ϕ_x_(t)* and *ϕ_y_(t)* of two given time series, *x(t)* and *y(t)*, the Hilbert transform (HT) is used to form the analytical signal *S(t)* as

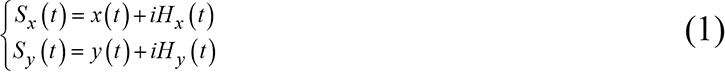

where and are the HT of two-time series, *x(t)* and *y(t)*, which are defined as,

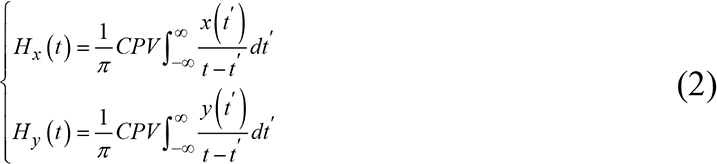

where the *CPV* denotes the Cauchy principal value. Afterward, corresponding analytical signal phases, *ϕ_x_(t)* and *ϕ_y_(t)*, can be computed as,

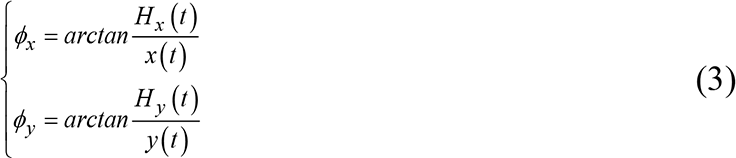

Finally, the PLV is formulated as

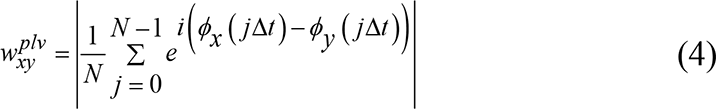

Where 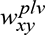 denotes the by PLV value between *x(t)* and *y(t)*, ^Δ^*^t^* denotes the sampling period, and *N* denotes the sample number.

#### Statistical analysis

Repeated-measure ANOVA with modality (auditory, visual, and audio-visual) as a within-participants variable and post-hoc tests (pairwise comparisons, Bonferroni-corrected) were used to quantify differences in ACC, RT, and power spectral strength. The assumption of sphericity was assessed with Mauchly’s test, and the Greenhouse-Geisser correction for non-sphericity was used to correct the *p*- values when required. Bonferroni correction was used for multiple pairwise comparisons. Topographical analyses were based on power spectral strength measured on 61 electrodes distributed over the entire scalp. Based on the constructed networks, network edges with significant differences (*p* < 0.01) among the three conditions were highlighted using paired *t*-tests. The mean amplitudes of P200 were analyzed using a two-way repeated-measure ANOVA with modality (3 levels) and electrode (6 levels) as within-participants variables. To estimate the statistical significance of differences between AV and sumAV, paired *t*-tests (two-tailed) was used.

### Experiment 2

#### Participants

The sample consisted of 15 healthy native speakers of Chinese (8 male, aged between 20 and 28, mean 23.03 years) from the students in UESTC. Experiments were conducted in accordance with the Declaration of Helsinki and within the current TMS safety guidelines of the International Federation of Clinical Neurophysiology [37]. None of the participants had a history of neurological or psychiatric disorders, and none of them was currently using any psychoactive medications. The experimental protocol was approved by the Institution Research Ethics Board of the UESTC. Written informed consent was obtained after the procedure had been fully explained, prior to scanning. All participants were paid Y 100 per hour for their participation.

#### Stimuli

The language-related stimuli in Experiment 2 were the same as those in Experiment 1.

#### Experimental procedures

We used a two × two × three factorial design with the three within-participants factors including TMS site (AF7 vs. FC5), TMS condition (effective vs. sham), and modality condition (auditory vs. visual vs. audio-visual). The experiment consisted of two site sessions (AF7 and FC5) that were performed with an inter-session interval of at least one week to avoid carry-over or earning effects, as depicts in Fig 3. Both sessions consisted of two blocks, separated by a 1 h break. The blocks differed with respect to the TMS condition (effective vs. sham). In each block, the tasks with stimuli in the three modalities (auditory, visual, and audio-visual) were performed in three runs, respectively. The participants were given a 3-minutes interval between two consecutive runs for rest, and the order of stimuli (i.e., auditory, visual, audio- visual) was randomized across participants. In each run, Chinese monosyllabic words were presented in sequences in random order or an order forming the two-phrase sentences for the three modalities. Each run consists of 25 trials including 20 standard trials and 5 outlier trials. The order of sentences and trials within a run was randomized. Participants were given a test to ensure that the rules of the task were understood before the experiment.

#### Continuous theta-burst stimulation

TMS was carried out with the Magstim Super Rapid 2 (The Magstim Company Ltd, UK), using a standard 70 mm figure-eight focal coil. The coil was placed tangentially to the skull with the coil handle oriented perpendicular to the target cortex, guided by the online BrainSight frameless stereotaxy system (BrainSight Frameless, Rogue Research, Montreal, Canada). Stimulation was delivered at an average of 40% (*SD* = 6.18) of the maximum stimulator output. No arms or other movements were elicited by the stimulation of the targeted site. Each ‘burst’ comprised three 50 Hz pulses and was repeated every 200 ms resulting in the delivery of 600 pulses over a 40 s period [38]. The sham stimulation was delivered using the cTBS protocol with the coil positioned at a perpendicular angle to AF7 or FC5 in a counterbalanced manner across participants. After cTBS or sham stimuli, participants were asked to perform the task and EEG and behavioral responses were simultaneously recorded.

#### Statistical analysis

RT, ACC, and P200 amplitudes were respectively subjected to three-way repeated-measures ANOVAs with TMS sites (2 levels), TMS condition (2 levels), and modality condition (3 levels) as within-participants variables and post-hoc tests were used to quantify differences between conditions. The power spectrum was analyzed using four-way repeated-measures ANOVAs with TMS sites (2 levels), TMS condition (2 levels), modality condition (3 levels), and power spectrum frequency (3 levels) as within-participants variables and post-hoc tests were used to quantify differences. The assumption of sphericity was assessed with Mauchly’s test, and the Greenhouse-Geisser correction for non-sphericity was used to correct the *p*- values when required. Bonferroni correction was used for multiple pairwise comparisons. Based on our hypothesis, we focused on interactions between TMS and modality conditions.

## Acknowledgments

We wish to thank all the participants who participated in Experiment 1 and 2.

## Supporting information

**S1 Fig.**
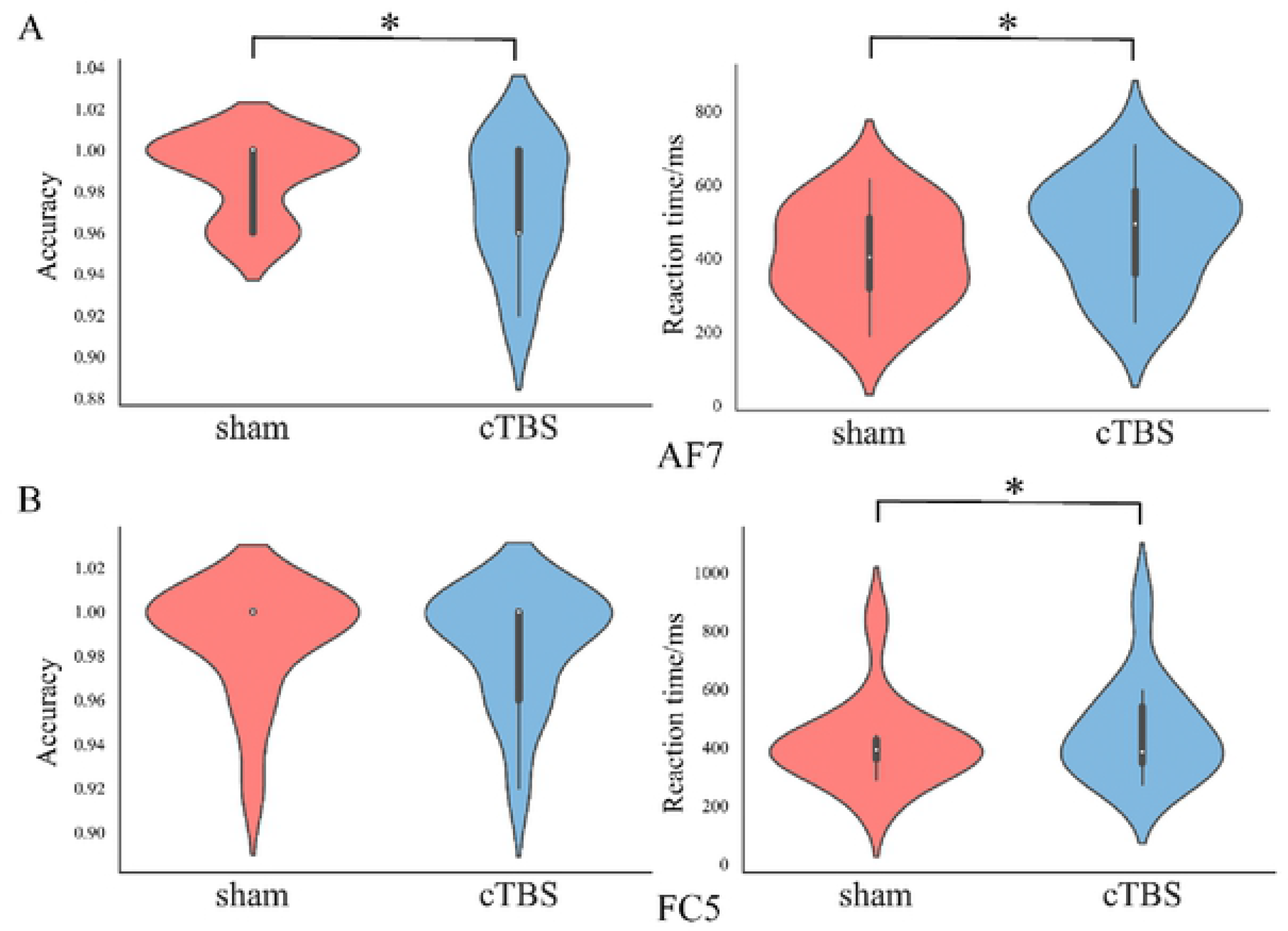
The mean ACC and RT in response to stimuli in the audio-visual modality following TMS stimulus and sham stimulation, shown in the form of violin plots. (A) cTBS and sham stimulation on AF7. (B) cTBS and sham stimulation on FC5. White circles represent the median, and the first and third quartiles are identified by the bottom and top of the bold vertical lines, respectively. The bottom and top of the thin vertical line represent the lower and upper adjacent values, respectively (* p < 0.05, Bonferroni-corrected).

**S2 Fig.**
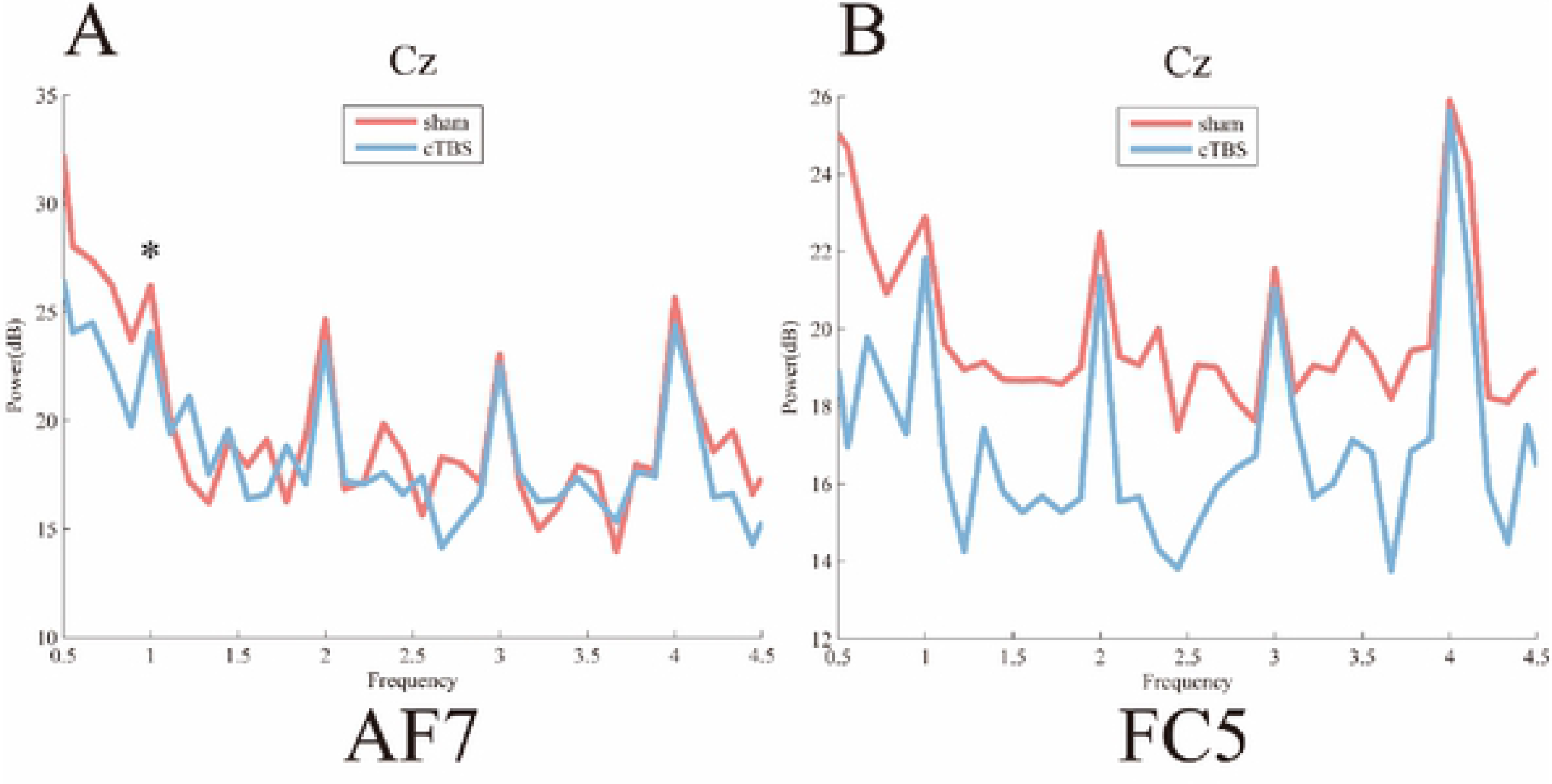
Power spectrum of audio-visual modality for each TMS condition. (A) cTBS on AF7 and sham stimulation. (B) cTBS on FC5 and sham stimulation. Error bars represent the standard errors of the mean acceptance rates. * *p* < 0.05.

**S3 Fig.**
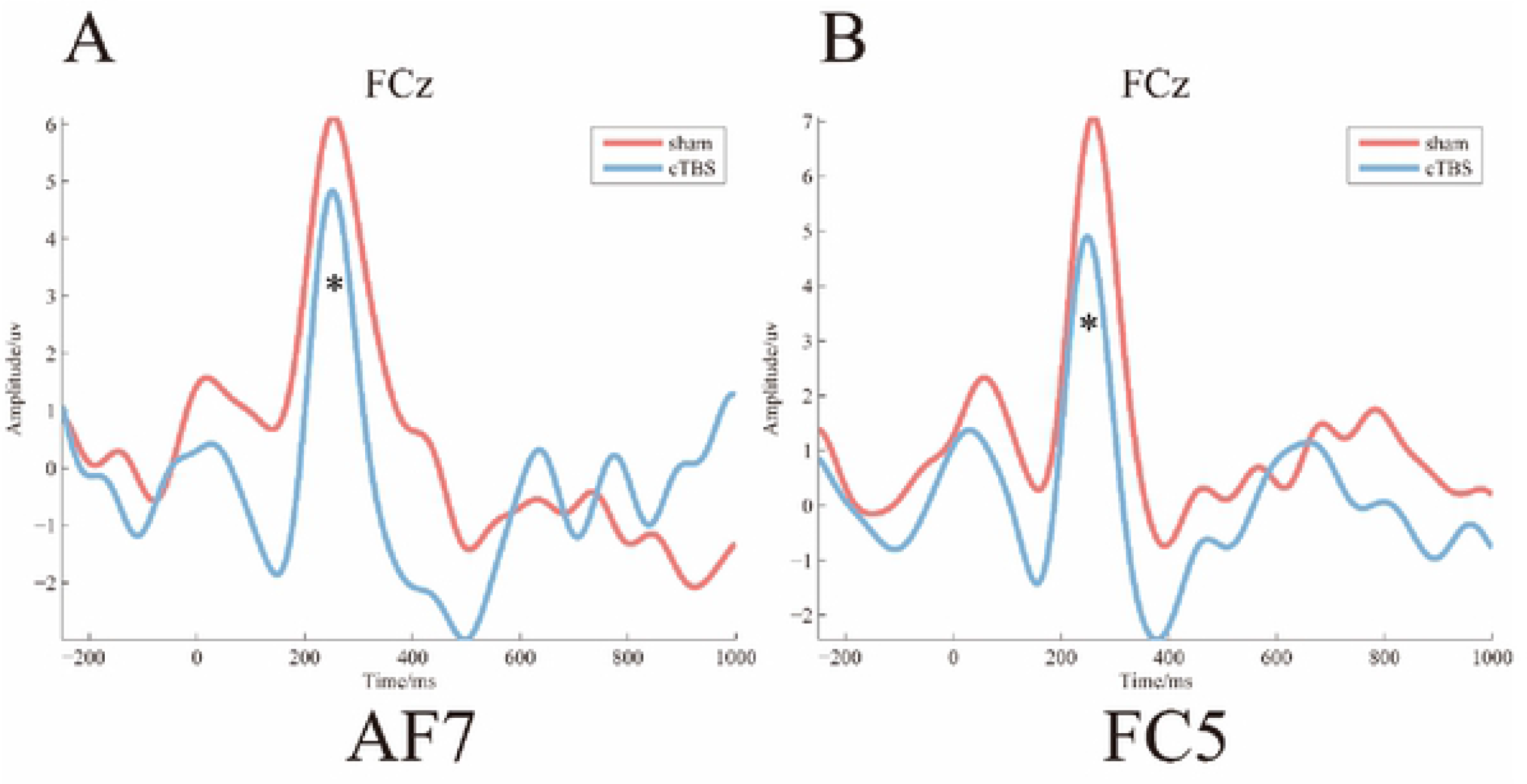
P200 of audio-visual condition for each TMS stimulus. (A) cTBS and sham stimulation on AF7. (B) cTBS and sham stimulation on FC5. Error bars represent the standard errors of the mean acceptance rates. * *p* < 0.05.

## References

1. Hagoort P. The neurobiology of language beyond single-word processing. Science. 2019;366(6461):55-+.

2. Jarvis ED. Evolution of vocal learning and spoken language. Science. 2019;366(6461):50-+.

3. Murray MM, Lewkowicz DJ, Amedi A, Wallace MT. Multisensory Processes: A Balancing Act across the Lifespan. Trends Neurosci. 2016;39(8):567–79.

4. Wallace MT, Woynaroski TG, Stevenson RA. Multisensory Integration as a Window into Orderly and Disrupted Cognition and Communication. Annual Review of Psychology, Vol 71. 2020;71:193–219.

5. Shao JJ. Xian, Dai Han Yu Tong Lun (The second edition): Shanghai Press; 2007.

6. Huang BR, Liao XD. Xian Dai Han Yu (Five enlarged edition): Higher Education Press; 2011.

7. van Atteveldt N, Formisano E, Goebel R, Blomert L. Integration of letters and speech sounds in the human brain. Neuron. 2004;43(2):271–82.

8. Basirat A, Brunelliere A, Hartsuiker R. The Role of Audio-visual Speech in the Early Stages of Lexical Processing as Revealed by the ERP Word Repetition Effect. Lang Learn. 2018;68:80–101.

9. Xu WY, Kolozsvari OB, Oostenveld R, Hamalainen JA. Rapid changes in brain activity during learning of grapheme-phoneme associations in adults. Neuroimage. 2020;220.

10. Arnal LH, Morillon B, Kell CA, Giraud AL. Dual Neural Routing of Visual Facilitation in Speech Processing. J Neurosci. 2009;29(43):13445–53.

11. Blomert L. The neural signature of orthographic-phonological binding in successful and failing reading development. Neuroimage. 2011;57(3):695–703.

12. Raij T, Uutela K, Hari R. Audio-visual integration of letters in the human brain. Neuron. 2000;28(2):617–25.

13. Ding N, Melloni L, Zhang H, Tian X, Poeppel D. Cortical tracking of hierarchical linguistic structures in connected speech. Nat Neurosci. 2016;19(1):158–64.

14. Berwick RC, Friederici AD, Chomsky N, Bolhuis JJ. Evolution, brain, and the nature of language. Trends Cogn Sci. 2013;17(2):89–98.

15. Luo H, Poeppel D. Phase patterns of neuronal responses reliably discriminate speech in human auditory cortex. Neuron. 2007;54(6):1001–10.

16. Ding N, Simon JZ. Emergence of neural encoding of auditory objects while listening to competing speakers. Proc Natl Acad Sci U S A. 2012;109(29):11854–9.

17. Zion Golumbic EM, Ding N, Bickel S, Lakatos P, Schevon CA, McKhann GM, et al. Mechanisms Underlying Selective Neuronal Tracking of Attended Speech at a “Cocktail Party”. Neuron. 2012;77(5):980–91.

18. Peelle JE, Gross J, Davis MH. Phase-Locked Responses to Speech in Human Auditory Cortex are Enhanced During Comprehension. Cereb Cortex. 2013;23(6):1378–87.

19. Pasley BN, David SV, Mesgarani N, Flinker A, Shamma SA, Crone NE, et al. Reconstructing Speech from Human Auditory Cortex. PLoS Biol. 2012;10(1).

20. Sheng JW, Zheng L, Lyu BJ, Cen ZH, Qin L, Tan LH, et al. The Cortical Maps of Hierarchical Linguistic Structures during Speech Perception. Cereb Cortex. 2019;29(8):3232–40.

21. Bemis DK, Pylkkanen L. Basic Linguistic Composition Recruits the Left Anterior Temporal Lobe and Left Angular Gyrus During Both Listening and Reading. Cereb Cortex. 2013;23(8):1859–73.

22. Giraud AL, Poeppel D. Cortical oscillations and speech processing: emerging computational principles and operations. Nat Neurosci. 2012;15(4):511–7.

23. Pallier C, Devauchelle AD, Dehaene S. Cortical representation of the constituent structure of sentences. Proc Natl Acad Sci U S A. 2011;108(6):2522–7.

24. Lerner Y, Honey CJ, Silbert LJ, Hasson U. Topographic Mapping of a Hierarchy of Temporal Receptive Windows Using a Narrated Story. J Neurosci. 2011;31(8):2906–15.

25. Si YJ, Wu X, Li FL, Zhang LY, Duan KY, Li PY, et al. Different Decision-Making Responses Occupy Different Brain Networks for Information Processing: A Study Based on EEG and TMS. Cereb Cortex. 2019;29(10):4119–29.

26. Li FL, Chen B, Li H, Zhang T, Wang F, Jiang Y, et al. The Time-Varying Networks in P300: A Task-Evoked EEG Study. IEEE Trans Neural Syst Rehabil Eng. 2016;24(7):725–33.

27. Ding N, Patel AD, Chen L, Butler H, Luo C, Poeppel D. Temporal modulations in speech and music. Neurosci Biobehav Rev. 2017;81(Pt B):181–7.

28. Dong L, Li FL, Liu Q, Wen X, Lai YX, Xu P, et al. MATLAB Toolboxes for Reference Electrode Standardization Technique (REST) of Scalp EEG. Front Neurosci. 2017;11.

29. Yao DZ. A method to standardize a reference of scalp EEG recordings to a point at infinity. Physiol Meas. 2001;22(4):693–711.

30. Picton TW, Bentin S, Berg P, Donchin E, Hillyard SA, Johnson R, et al. Guidelines for using human event-related potentials to study cognition: Recording standards and publication criteria. Psychophysiology. 2000;37(2):127–52.

31. Luck SJ. An introduction to the event-related potential technique. Cambridge, MA: MIT press; 2014.

32. Christoffels I, Timmer K, Ganushchak L, La Heij W. On the production of interlingual homophones: delayed naming and increased N400. Lang Cogn Neurosci. 2016;31(5):628–38.

33. Sakkalis V. Review of advanced techniques for the estimation of brain connectivity measured with EEG/MEG. Comput Biol Med. 2011;41(12):1110–7.

34. Sun J, Li Z, Tong S. Inferring functional neural connectivity with phase synchronization analysis: a review of methodology. Comput Math Methods Med. 2012;2012:239210.

35. Srinivasan R, Nunez PL, Silberstein RB. Spatial filtering and neocortical dynamics: Estimates of EEG coherence. IEEE Trans Biomed Eng. 1998;45(7):814–26.

36. Xu P, Xiong XC, Xue Q, Li PY, Zhang R, Wang ZY, et al. Differentiating Between Psychogenic Nonepileptic Seizures and Epilepsy Based on Common Spatial Pattern of Weighted EEG Resting Networks. IEEE Trans Biomed Eng. 2014;61(6):1747–55.

37. Rossi S, Hallett M, Rossini PM, Pascual-Leone A, Avanzini G, Bestmann S, et al. Safety, ethical considerations, and application guidelines for the use of transcranial magnetic stimulation in clinical practice and research. Clin Neurophysiol. 2009;120(12):2008–39.

38. Huang YZ, Edwards MJ, Rounis E, Bhatia KP, Rothwell JC. Theta burst stimulation of the human motor cortex. Neuron. 2005;45(2):201–6.

39. Weintraub-Brevda RR, Chua EF. The role of the ventrolateral prefrontal cortex in emotional enhancement of memory: A TMS study. Cogn Neurosci. 2018;9(3-4):116–26.

40. Scrivener CL, Reader AT. Variability of EEG electrode positions and their underlying brain regions: visualising gel artifacts from a simultaneous EEG-fMRI dataset. 2021.

41. Gao S, Zika O, Rogers RD, Thierry G. Second Language Feedback Abolishes the “Hot Hand” Effect during Even-Probability Gambling. J Neurosci. 2015;35(15):5983–9.

42. Mittag M, Alho K, Takegata R, Makkonen T, Kujala T. Audio-visual attention boosts letter-speech sound integration. Psychophysiology. 2013;50(10):1034–44.

43. Bemis DK, Pylkkanen L. Simple Composition: A Magnetoencephalography Investigation into the Comprehension of Minimal Linguistic Phrases. J Neurosci. 2011;31(8):2801–14.

44. Senkowski D, Saint-Amour D, Gruber T, Foxe JJ. Look who’s talking: The deployment of visuo-spatial attention during multisensory speech processing under noisy environmental conditions. Neuroimage. 2008;43(2):379–87.

45. Baier B, Kleinschmidt A, Muller NG. Cross-modal processing in early visual and auditory cortices depends on expected statistical relationship of multisensory information. J Neurosci. 2006;26(47):12260–5.

46. Bushara KO, Hanakawa T, Immisch I, Toma K, Kansaku K, Hallett M. Neural correlates of cross-modal binding. Nat Neurosci. 2003;6(2):190–5.

47. Romei V, Murray MM, Merabet LB, Thut G. Occipital Transcranial magnetic stimulation has opposing effects on visual and auditory stimulus detection: Implications for multisensory interactions. J Neurosci. 2007;27(43):11465–72.

48. van Atteveldt NM, Formisano E, Blomert L, Goebel R. The effect of temporal asynchrony on the multisensory integration of letters and speech sounds. Cereb Cortex. 2007;17(4):962–74.

49. Grodzinsky Y, Santi A. The battle for Broca’s region. Trends Cogn Sci. 2008;12(12):474–80.

50. Friederici AD. The Brain Basis of Language Processing: From Structure To Function. Physiol Rev. 2011;91(4):1357–92.

51. Matchin W, Hammerly C, Lau E. The role of the IFG and pSTS in syntactic prediction: Evidence from a parametric study of hierarchical structure in fMRI. Cortex. 2017;88:106–23.

52. Hickok G, Poeppel D. Opinion - The cortical organization of speech processing. Nature Reviews Neuroscience. 2007;8(5):393–402.

53. Singhal A, Doerfling P, Fowler B. Effects of a dual task on the N100-P200 complex and the early and late Nd attention waveforms. Psychophysiology. 2002;39(2):236–45.

54. Ghani U, Signal N, Niazi IK, Taylor D. ERP based measures of cognitive workload: A review. Neurosci Biobehav Rev. 2020;118:18–26.

55. Zou Y, Tsang YK, Wu Y. Semantic Radical Activation in Chinese Phonogram Recognition: Evidence from Event-Related Potential Recording. Neuroscience. 2019;417:24–34.

56. Tong XH, Wang Y, Tong SX. Neurocognitive Correlates of Statistical Learning of Orthographic-Semantic Connections in Chinese Adult Learners. Neurosci Bull. 2020;36(8):895–906.

57. Kiyonaga K, Grainger J, Midgley K, Holcomb PJ. Masked cross-modal repetition priming: An event-related potential investigation. Lang Cogn Process. 2007;22(3):337–76.

58. Kutas M, Federmeier KD. Thirty Years and Counting: Finding Meaning in the N400 Component of the Event-Related Brain Potential (ERP). Annu Rev Psychol. 2011;62:621–47.

59. Coltheart M, Rastle K, Perry C, Langdon R, Ziegler J. DRC: A dual route cascaded model of visual word recognition and reading aloud. Psychol Rev. 2001;108(1):204–56.

60. Bergelson E, Swingley D. At 6-9 months, human infants know the meanings of many common nouns. Proc Natl Acad Sci U S A. 2012;109(9):3253–8.

61. Scott SK. From speech and talkers to the social world: The neural processing of human spoken language. Science. 2019;366(6461):58–61.

